# Glial voltage-gated K^+^ channels modulate the neural abiotic stress tolerance of *Drosophila melanogaster*

**DOI:** 10.1101/2024.06.28.601156

**Authors:** Mads Kuhlmann Andersen, Dawson B. H. Livingston, R. Meldrum Robertson, Heath A. MacMillan

## Abstract

Severe abiotic stress causes insects to lose nervous function and enter a state of paralytic coma. Central to this loss of function is a spreading depolarization (SD), where a characteristic collapse of ion gradients depolarizes neuronal and glial membranes and rapidly shuts down the CNS. Despite representing a critical limit to CNS function, the stress threshold that elicits SD can be altered by the process of acclimation, though the mechanisms underlying this response remain largely unknown. Here, we made electrophysiological measurements of SD and investigated the role of K^+^ channels in acclimation of the CNS stress response of *Drosophila melanogaster*. First, we demonstrate that improved cold tolerance in the CNS elicited by cold acclimation was abolished by pharmacological blockade of K^+^ channels with voltage-gated K^+^ channels representing most of this effect. Next, we used the UAS/Gal4 model system to screen for candidate genes encoding glial voltage-gated K^+^ channels and found that knockdown of *sei*- and *Shaw*-encoded channels mimicked the effect of K^+^ blockade in cold-acclimated flies. Furthermore we show that the knockdown of glial *sei*-encoded channels also impair tolerance to anoxia and heat stress. These findings suggest that voltage-gated K^+^ channels, especially those encoded by *sei*, are integral to the CNS stress- and acclimation-response and we posit that this is elicited through mechanisms involving glial spatial buffering and barrier function. Establishing such causal links between tissue-specific expression of candidate genes and SD mechanisms will inevitably aid our understanding of insect ecophysiology and SD-related neuropathologies.

**New and Noteworthy:** Using thermal acclimation and pharmacology, we demonstrate that voltage-gated K^+^ channels are involved in setting the threshold for cold-induced spreading depolarization (SD) in the *Drosophila melanogaster* CNS. Glial knockdown of channels encoded by *sei* and *Shaw* reduced the resistance to cold-induced SD, highlighting their importance in acclimation of the CNS. Glia-specific *sei*-knockdown also reduced tolerance to anoxia and heat. We posit that *sei*-channels are involved the CNS stress- and acclimation-responses through glial spatial buffering mechanisms.

## Introduction

Abiotic stressors pose a critical challenge for the fitness and survival of insects because they directly affect molecular, cellular, and physiological functions (1–3). If sufficiently stressed, insects will eventually enter a comatose state referred to as chill-, heat-, or anoxic coma depending on the stressor experienced (4–6). These comas are characterized by complete unresponsiveness and inability to move, implicating the neuromuscular system (7, 8). Indeed, the loss of equilibrium prior to the comatose state and subsequent complete loss of organismal function has been linked to a rapid loss of central nervous system (CNS) function (9, 10). The phenomenon leading to the abrupt arrest of neural function is known as spreading depolarization (SD). This same event occurs at the critical thermal limits (e.g. CT_max_ and CT_min_) in insects (11–14) and rapidly shuts down the CNS during anoxia exposure (15–17).

During SD, ion gradients across neural and glial cell membranes collapse in an all-or-nothing event that leads to an almost complete depolarization of neuronal membrane potentials (9, 18). This debilitating event then spreads into adjacent neural tissue, resulting in a wave of neural depolarization that incapacitates the entire neuropil, but is also confined to it as it cannot spread into neural connectives (16). Spreading depolarization was first discovered in the white matter of mammals eight decades ago (19) and has since been linked to several human neuropathologies (e.g. migraine, stroke, traumatic injury) (20, 21). The mechanisms underlying SD initiation and propagation, however, remain unknown (18, 22–24). This event was first observed in the insect brain (cerebral ganglion) two decades ago (25, 26) and the phenomenology of SD appears remarkably similar in insects and mammals (9). Consequently, insects have now been established as valuable model systems for SD research (9, 15, 16), and fruit flies are in particular being used as translational models for stroke (27) and seizure disorders (28, 29). Interestingly, SD has also been observed in the CNS in fish at the CT_max_ (30). Thus, developing our understanding of SD in insects can not only reveal the physiological processes limiting abiotic stress tolerance of ectothermic animals or promoting variation therein, but can also provide valuable insights into the mechanisms of SD initiation, propagation, and susceptibility in humans and other mammals.

While SD represents a clear limit to central nervous system function, the dose of abiotic stress needed to trigger SD is highly plastic and depends on multiple factors; one of the most studied being thermal history. Most insects are able to alter their stress tolerance through acclimation, seasonal acclimatization, or brief thermal pre-treatments (e.g. hardening) (31). For example, the fruit fly (*Drosophila melanogaster*) has an impressive capacity to improve its resistance to cold-induced SD in response to cold acclimation or rapid hold hardening (32–34), and this can result in ∼ 10°C of difference in the CT_min_ among flies experiencing these treatments (35–37). Similarly, differences in critical thermal limits among *Drosophila* species have been leveraged to show that these limits are strong predictors of species distribution (38–43). As SD underlies the organismal loss of function at the critical thermal limits, understanding the mechanisms underlying induced variation in SD in *Drosophila* and other insects will aid our understanding of the molecular, cellular, and physiological processes ultimately determining their thermal niches.

Despite the precise triggering mechanism of SD being unknown, recent research has highlighted a wide range of processes involved in modulating the susceptibility and temperature threshold for SD (see (10) and (44)). These processes include improved maintenance of Na^+^/K^+^- ATPase activity at low temperature (32, 45, 46), increased abundance of Na^+^:K^+^:2Cl^-^ symporters and K^+^ channels (47), and maintained octopaminergic regulation of K^+^ conductances in perineurial glia (48). These changes agree with the general view that improvements of the capacity to regulate osmotic and ionic gradients are key adaptations to low temperature in most terrestrial insects (8, 49–51). The basic components and mechanisms of ion regulation within the CNS of insects and mammals are remarkably similar, and consist of specialized glial cells acting to counteract the changes in Na^+^ and K^+^ gradients elicited by neural activity (52–54). Of particular importance here is the process of glial spatial buffering, by which elevations in the extracellular concentration of K^+^ within the CNS (i.e. interstitial K^+^ concentration) are counteracted by the uptake and shuttling of K^+^away from the interstitium through the electrically coupled glia (55–58). Glial spatial buffering is maintained by a complex combination of processes, but as expected the Na^+^/K^+^-ATPase and K^+^ channels play key roles as active and passive components, respectively (56, 58). During SD, the interstitial K^+^ concentration increases rapidly and its diffusion has been proposed as a mechanism for SD propagation (22, 59, 60). Thus, it is not surprising that K^+^ channels have been highlighted as modulators of the stress threshold for induction and propagation of SD (16, 47, 61–64), as well as a number of SD-associated neuropathological phenotypes, like epilepsy and seizure, in insects and mammals alike (65–68). A recent study highlighted the role of glia- and neuron-specific expression of voltage-gated K^+^ channels in the response to heat stress and by showing clear reductions in organismal thermal tolerance of *Drosophila melanogaster* (i.e. time to paralysis/coma) after transgenic knockdown of several channel-encoding genes (69). In this case, the findings were not tied to a specific physiological phenomenon, but we speculate that the loss of K^+^ channels likely modified either neural excitability or glial spatial buffering, which would elicit increased susceptibility to SD and lead to a more temperature-sensitive phenotype.

In the present study, we aimed to test the hypothesis that a reduction in expression of genes encoding voltage-gated K^+^ channels can modify organismal thermal limits by increasing susceptibility to SD. First, we investigated the role of voltage-gated K^+^ channels in the thermal acclimation response of *D. melanogaster*. We took an electrophysiological approach to monitor the loss of CNS function by SD and compared the effects of K^+^ channels blockers on the low-temperature threshold for SD in both cold- and warm-acclimated flies. Cold-acclimation greatly improved the resistance to cold-induced SD, but this effect was strongly blunted by application of a voltage-gated K^+^ channel blocker. Next, we used the UAS/Gal4 model system to produce tissue-specific knockdowns of voltage-gated channel-encoding genes and screen for candidate genes involved in setting the threshold for cold-induced SD. Here, glia were targeted for the tissue-specific knockdown for multiple reasons: 1) Glial, not neural, expression of heat shock proteins (which are integral to the thermal stress response) protects K^+^ homeostasis of the brain when stressed (70), 2) glial AMPK activation, not neuronal, is confers neuroprotection following anoxic SD (71), and 3) recent evidence suggests that SD is triggered at the glial hemolymph-brain barrier in insects (72). We found that the glial knockdown of *sei* elicited the most pronounced effects, and we therefore tested the effects of knockdown of this gene on anoxia-induced SD and recovery.

Lastly, we tested how the *sei* knockdown affected organismal heat tolerance. Because we used the same crosses as Hill et al. (69) for this experiment, it also served to test the repeatability of glial RNAi-knockdown of *sei* using the UAS/Gal4 model system.

## Materials and Methods

### Fly lines and husbandry

For experiments involving acclimation and/or pharmacology, we used a line of *Drosophila melanogaster* (Meigen 1830) established from isofemales from a population collected near London and Niagara on the Lake in Ontario, Canada (73). These flies were kept in an incubator (MIR-154- PA, Panasonic) at 20°C in 200 mL bottles with ∼ 40 mL banana-based diet at the bottom (recipe in (74)), and with a 12:12 light:dark cycle.

For transgenic experiments, our background Canton S line and glial-targeting *repo*-Gal4 line(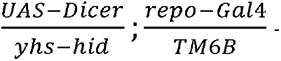– based on the Canton S background line) were obtained from the lab of Dr. Woo-Jae Kim (University of Ottawa). UAS-RNAi TRiP (Transgenic RNAi Project, see (75)) lines were obtained from the Bloomington Drosophila Stock Centre and were as follows (full gene name and stock # in parentheses): *eag* (*ether-a-go-go*, #31678), *sei* (*seizure*, #31861), *Sh* (*shaker*, #53347), *Shab* (Shaker cognate b, #25805), *Shal* (Shaker cognate l, #31879), and *Shaw* (Shaker cognate w, #28346). These populations were kept in 200 mL bottles with ∼ 40 mL of standard Bloomington medium with a 12:12 light cycle at 25°C, except the *repo*-Gal4 line which was kept at 18°C (also in MIR-154-PA incubators).

### Fly preparation and electrophysiology

Individual flies were held by the head (using gentle aspiration and a 100 μL pipette tip) and fixed in a thin layer of wax on top of a glass cover slide using micro-forceps. This procedure eliminated the need for CO_2_ or other anaesthetics immediately prior to experiments, which would have otherwise confounded our results (76, 77). From here, two holes were cut in the cuticle of the fly with a pair of micro-scissors: 1) immediately above the ocelli on the head to access the brain, and 2) towards the end of the abdomen to gain access to the hemolymph for electrical grounding.

The loss of central nervous system function by spreading depolarization was observed electrophysiologically following previously described methods (78). For these observations we used glass microelectrodes fashioned from filamented borosilicate glass (1B100F-4, World Precision Instruments (WPI), Sarasota, Fl, USA) and pulled to a tip resistance of 5-7 MΩ in a Flaming-Brown P1000 electrode puller (Sutter Instruments, Novato, CA, USA). Glass microelectrodes were backfilled with 500 mM KCl, placed in an electrode holder with a Ag/AgCl wire, attached to a micromanipulator (M3301-M3-L, WPI), and connected to a Duo 773 intracellular electrometer (WPI). Raw outputs from the electrometer were digitized with a PowerLab 4SP A/D converter (ADInstruments, Colorado Springs, CO, USA) and fed to a computer running LabChart 4.0 software (ADInstruments).

Once prepared, the cover slide with the fly affixed was quickly moved to a custom-built thermoelectric (Peltier-style) cooling stage under the microscope. Here, a type K thermocouple was placed immediately next to the fly’s head to monitor the head and brain temperature. Temperatures measured by the thermocouple were fed through a TAC80B-K thermocouple-to-analog converter (Omega, Stamford, CT, USA) and the PowerLab A/D converter, and read by the LabChart software to obtain concurrent measurements of temperature and electrical potential. Next, an Ag/AgCl wire was inserted into the hole in the abdomen to ground the preparation, after which the micromanipulator was used to insert the electrode into the brain from a dorsolateral position.

Successful placement of the electrode was indicated by a sudden, small (5-10 mV) increase in the measured potential, which is indicative of having penetrated the hemolymph-brain barrier and represents the transperineurial potential (TPP) (9, 74). From here, temperature was lowered by 1°C min^-1^ until an abrupt drop in the TPP was observed (∼ 30-50 mV in 5-10 s), which indicates that a spreading depolarization has occurred (9, 78), and the temperature of onset was quantified as the temperature at the half-amplitude of the drop in TPP (see **Fig. 1**).

**Figure 1.**
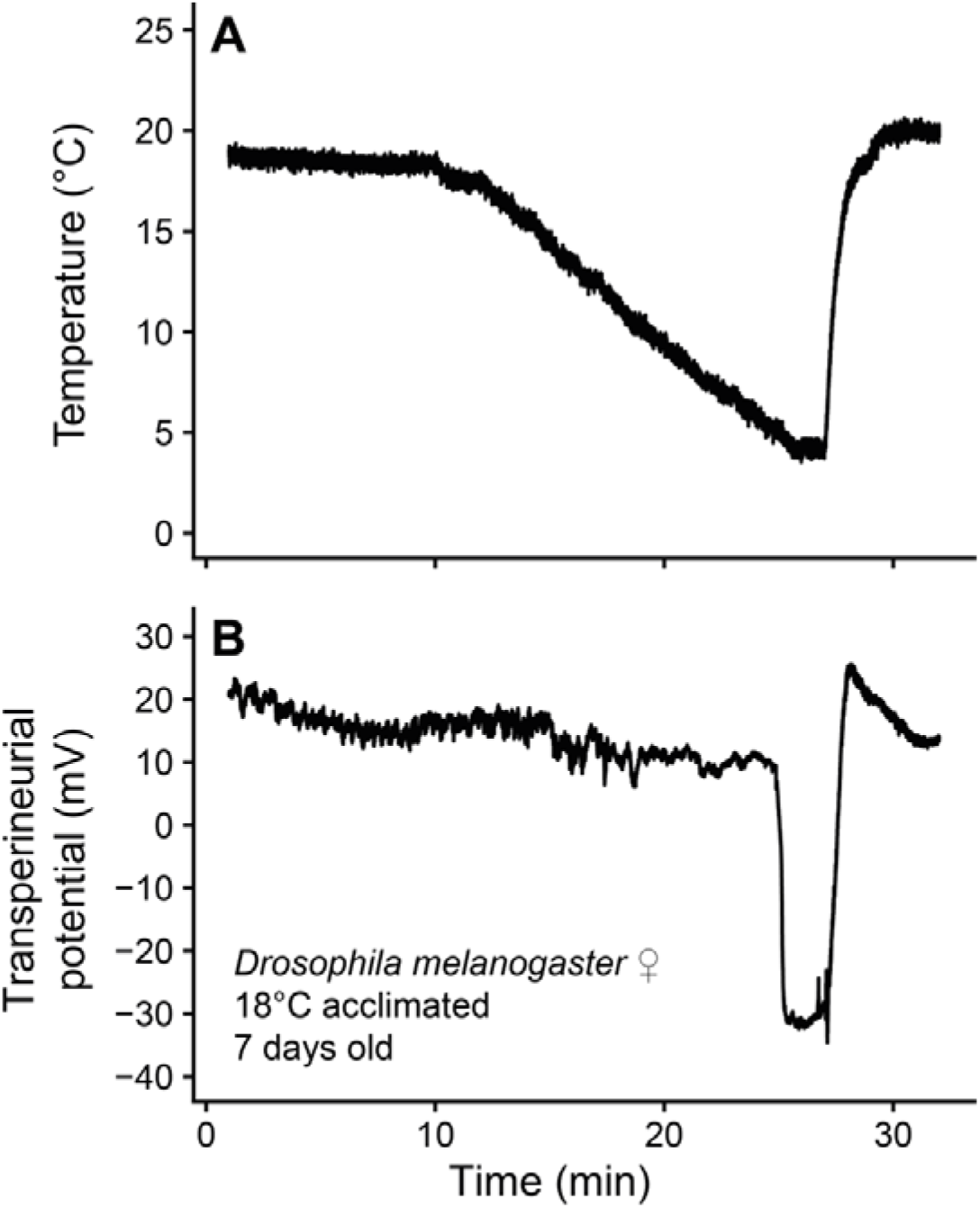
– Cold-induced spreading depolarization in *Drosophila melanogaster*. After a waiting period, temperature (A) was lowered by 1°C min^-1^ during which the TPP (B) was continuously monitored. At a critical temperature, the TPP exhibits a rapid negative shift, which denotes the onset of SD. The temperature at the half-amplitude of the negative shift is noted as the temperature leading to cold-induced SD; in this case ∼ 5°C.

### Experiment 1 – Effects of acclimation and pharmacological interventions on the spreading depolarization temperature

Experimental flies were produced by having parental flies from the 20°C population oviposit in bottles with fresh medium for ∼ 4 h, after which they were moved back into their rearing bottles and the now egg-containing bottles were moved into incubators to acclimate at either 18°C or 25°C with the same 12:12 light cycle. Upon emergence, developmentally acclimated flies were tipped into 40 mL vials with ∼ 7 mL of medium and left to mature at their respective acclimation temperature for seven days prior to experiments. Only females were used in experiments.

On the day of the experiment, flies from either acclimation group were selected for one of four treatments: 1) control, 2) sham, 3) voltage-gated K^+^ channel block, or 4) general K^+^ channel block. Control flies were prepared for electrophysiological experiments as described above and after a 10 min wait at room temperature they were cooled from room temperature (22-23°C) at 1°C min^-1^ until a spreading depolarization was observed and its temperature recorded. Sham flies were prepared similarly, except they had 14-15 nL saline (147 mM NaCl, 10 mM KCl, 4 mM CaCl_2_, and 10 mM HEPES; pH 7.2) injected into the head capsule, through the hole cut for the electrode and after electrode insertion, with a Nanoject II Injector (Drummond Scientific Company, Broomall, PA, USA) before the 10 min wait. For groups 3 and 4, flies were prepared in the same way as the sham group, except the saline contained an additional 1 mM of 4-aminopyridine (4-AP, a voltage- gated K^+^ channel blocker, Sigma-Aldrich, St Louis, MO, USA) or 1 mM tetraethylammonium chloride (TEA, a general K^+^ channel blocker, also Sigma-Aldrich) with NaCl being lowered accordingly to maintain saline osmolality. Pilot experiments revealed that neither the sham, 4-AP, or TEA injections resulted in any spreading depolarization at room temperature within the ∼ 30 min timeline of our experiments. N = 12 flies per acclimation and treatment group for a total of 96 flies.

### Experiment 2 – Screen of voltage-gated K^+^ channel gene knockdowns

To generate experimental flies, bottles containing larvae were heated to 37°C for 90 min and after which they were moved to 25°C rearing conditions. This treatment kills or sterilizes all male flies in the *repo*-Gal4 line (due to *hs-hid* expression linked to the Y chromosome; *yhs-hid*) and ensures that emerging females are virgins before any crossing (79, 80). The other populations were given the same treatment to prevent potential transgenerational effects of the heat shock, and this had no apparent effect on viability. Flies emerging from the resulting pupae were removed from their bottles every hour and sorted under brief CO_2_ anaesthesia to ensure that females remained virgins. Males and females were moved to different 40 mL vials with ∼ 7 mL Bloomington medium and kept here for two days to remove potential effects of the CO_2_ anaesthesia (77). After recovering from CO_2_ anaesthesia, 10 females and 5 males were moved to the same 40 mL vials and left to mate and oviposit for two days, and after two more days in another fresh vial the adults were discarded. The specific crosses we used are outlined in **Table 1**. Upon emergence of the F1 generation, these flies were sorted daily under brief CO_2_ anaesthesia and females were left to mature until the experiments were carried out at seven days of age. On the day of the experiment, flies were prepared as described above and cooled from room temperature (22-23°C) at a rate of 1°C min^-1^ until a spreading depolarization was observed and its temperature recorded. N = 16 for each cross, except the *repo*-*Sh*^RNAi^ cross which had 15, for a total of 223 flies.

**Table 1.** – Transgenic crosses performed and used in experiments along with their sample sizes for each experiment.

| Parental crosses (♀ x ♂) | F<sub>1</sub> genotype (♀) | Group purpose | Experimental sample sizes |  |  |
| --- | --- | --- | --- | --- | --- |
|  |  |  | Exp. 2 | Exp. 3 | Exp. 4 |
| wt × wt | wt (+ > +) | Control (wildtype) | 16 | 16/15* | 9 |
| *repo*-Gal4 × wt | *repo* > + | Control (driver) | 16 | 16 | 11 |
| wt × UAS-TRiP-*eag* | + > *eag*<sup>RNAi</sup> | Control (reporter) | 16 | - | - |
| *repo*-Gal4 × UAS-TRiP-*eag* | *repo* > *eag*<sup>RNAi</sup> | RNAi knockdown | 16 | - | - |
| wt × UAS-TRiP-*sei* | + > *sei*<sup>RNAi</sup> | Control (reporter) | 16 | 16 | 12 |
| *repo*-Gal4 × UAS-TRiP-*sei* | *repo* > *sei*<sup>RNAi</sup> | RNAi knockdown | 16 | 16 | 12 |
| wt × UAS-TRiP-*Sh* | + > *Sh*<sup>RNAi</sup> | Control (reporter) | 15 | - | - |
| *repo*-Gal4 × UAS-TRiP-*Sh* | *repo* > *Sh*<sup>RNAi</sup> | RNAi knockdown | 16 | - | - |
| wt × UAS-TRiP-*Shab* | + > *Shab*<sup>RNAi</sup> | Control (reporter) | 16 | - | - |
| *repo*-Gal4 × UAS-TRiP-*Shab* | *repo* > *Shab*<sup>RNAi</sup> | RNAi knockdown | 16 | - | - |
| wt × UAS-TRiP-*Shal* | + > *Shal*<sup>RNAi</sup> | Control (reporter) | 16 | - | - |
| *repo*-Gal4 × UAS-TRiP-*Shal* | *repo* > *Shal*<sup>RNAi</sup> | RNAi knockdown | 16 | - | - |
| wt × UAS-TRiP-*Shaw* | + > *Shab*<sup>RNAi</sup> | Control (reporter) | 16 | - | - |
| *repo*-Gal4 × UAS-TRiP-*Shaw* | *repo* > *Shab*<sup>RNAi</sup> | RNAi knockdown | 16 | - | - |

\*One wt fly was removed from the analysis of recovery times in Exp. 3 as it had an unusually long recovery time (>3x longer than the wt flies).

### Experiment 3 – Effect of sei knockdown on anoxia resistance

Experimental flies were prepared as described for *Experiment 2* above, except only the *sei* gene was knocked down (see **Table 1**). Flies were prepares for electrophysiological experiments as described above, however, instead of cold they were exposed to anoxia. This experiment was set up in such a way that flies were continuously exposed to a flow of atmospheric air (created by a small air pump) which could be switched to pure, compressed N_2_ through a series of three-way valves (81). Gasses were humidified by bubbling them through separate flasks of room temperature water before they were delivered to the fly through a piece of Nalgene rubber tubing (1/8”, ThermoFischer Scientific, Waltham, Ma, USA) placed ∼ 1.5 cm laterally of fly. Gas flow was adjusted to ∼ 0.8 L min^-1^ using an adjustable tube clamp. From here, the TPP was monitored continuously while flies were exposed to anoxia for three minutes, after which the N_2_ was switched back to atmospheric air to observe the TPP recovery. The time to anoxic spreading depolarization was quantified as the time from the switch to anoxia to the half-amplitude of the drop in TPP (44), and the recovery time as the time to reach half-amplitude of the TPP recovery after reintroduction of oxygen (71) (see **Fig. 2**). N = 16 for each of the four crosses used here for a total of 64 flies (see **Table 1**).

**Figure 2.**
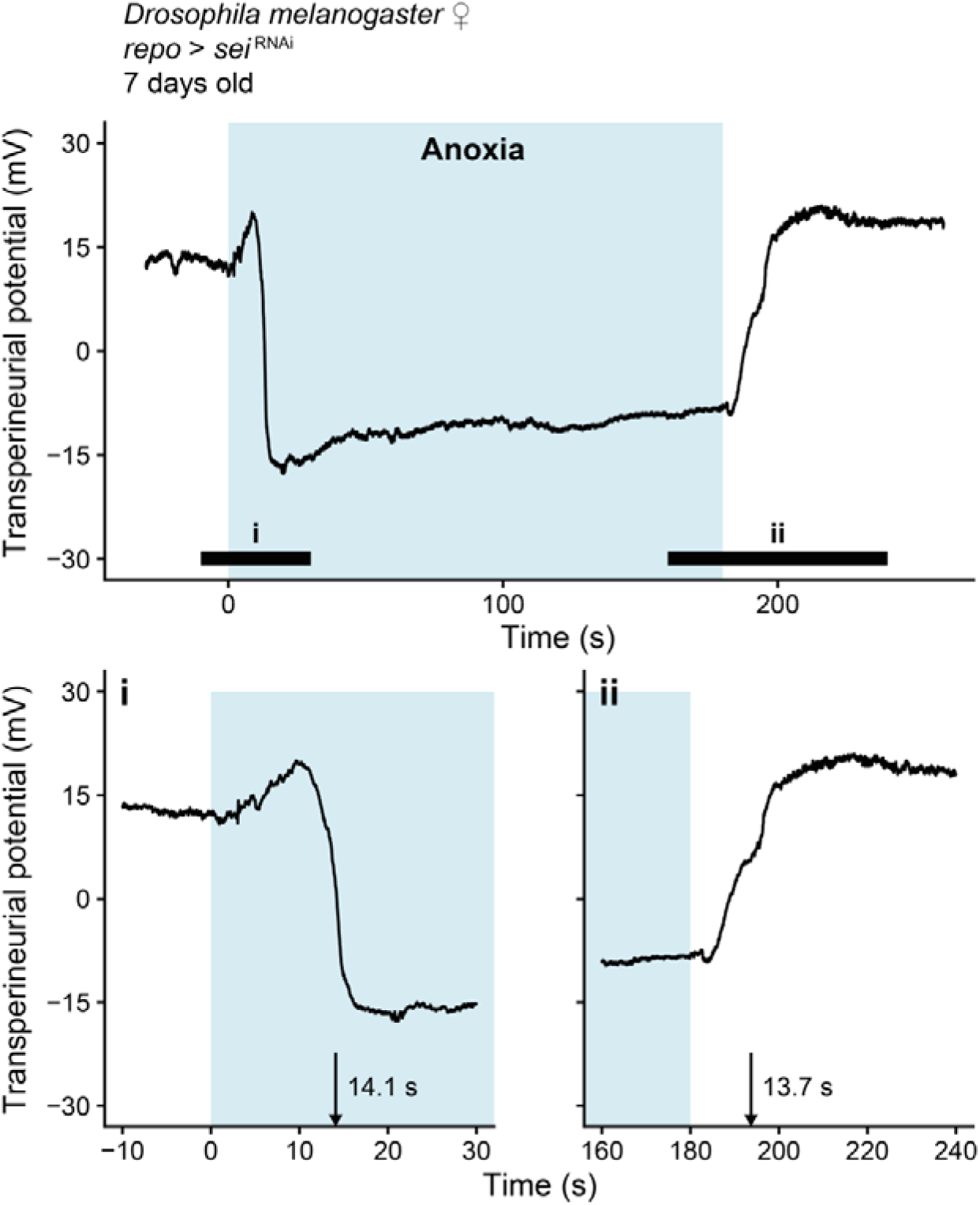
– Onset and recovery from anoxia-induced spreading depolarization in *Drosophila melanogaster*. The TPP is continuously monitored while an atmospheric air stream running over the fly is switched to pure N_2_ gas (shaded light blue area). This rapidly results in a negative shift in the TPP after a brief overshoot, signifying an anoxia-induced SD, where (i) the time to onset is noted as the time to reach the half-amplitude of the negative TPP shift. Following the return of oxygen (ii) the TPP exhibits a fast, positive shift and recovery to baseline, where the time to recover is noted as the time from the return of oxygen to the half-amplitude of the positive shift.

### Experiment 4 – Effect of sei knockdown on heat resistance

Flies for experiments were prepared as described for *Experiment 2*, but like *Experiment 3* only for the *sei* gene. Instead of electrophysiological experiments, these flies were used to observe the behavioural response to stressful heat following the approach outlined by Overgaard et al. (82).

This experiment was included to confirm that the phenotypes of the UAS-Gal4 crosses used here were similar to those obtained from similar crosses with a different background line (*w*^1118^, see Hill et al. (69)). Furthermore, heat exposure is a common way to induce SD in insects (16), and like the CT_min_, the CT_max_ also correlates strongly with timing of SD (11). Briefly, flies were placed in individual 4 mL glass vials and submerged into a water bath held at 40°C. From here, they were observed continuously, and the times until 1) loss of coordinated movements (i.e. loss of righting reflex) and 2) complete paralysis were recorded for each fly. Throughout the heat exposure, flies were encouraged to move by gently tapping the vials with a small metal rod. N = 9-12 with a total of 44 flies (see **Table 1** for more details).

### Data analysis

All statistical analyses were performed in R software (v. 4.3.2, R Core Team (83)). Normality of all datasets was confirmed with Shapiro-Wilks tests.

The isolated and interactive effects of acclimation temperature and pharmacological intervention (i.e. sham, 4-AP, and TEA) on the temperature that led to spreading depolarization were investigated with a two-way ANOVA. As a follow-up, we normalized the controls and K^+^ channel blocks to the sham group to directly compare the effects of the interventions to the controls in each acclimation group. This dataset (now without the sham group) was analyzed using a two-way ANOVA on log-transformed data.

The RNAi knockdown of each gene was compared to three controls with a one-way ANOVA: 1) the wildtype control, 2) the driver control, and 3) its gene-specific reporter control. If an effect on the spreading depolarization temperature was found, this was followed by Dunnett’s test, which compares a single group to multiple other groups without making comparisons among the multiple other groups. Thus, only if the knockdown group was statistically different from all three controls was it deemed to have a significant effect. These tests were performed using the DunnettTest() function from the ‘*DescTools*’ package (84). A similar analysis was performed on the effects of the *sei* knockdown on anoxia-induced spreading depolarization and recovery, and the organismal heat tolerance. The outputs of these analyses can be found in **Tables S1** and **S2**. Note that one extreme outlier was removed from the anoxia recovery dataset as it recovered more than three times slower than others from the same group (wild type); its’ time to anoxia value was within the normal range.

In all analyses the level of statistical significance was 0.05. Values reported represent means ± their standard error unless otherwise stated.

## Results

### The role of K^+^ channels in the acclimation response

To test whether voltage-gated K^+^ channels are involved in setting the limit for cold-induced spreading depolarization, we induced variation in the threshold temperature with thermal acclimation and subsequently tested whether the application of a sham treatment (saline) or a pharmacological blockade of either voltage-gated K^+^ channels (with 4-AP) or K^+^ channels in general (with TEA) increased the temperature leading to spreading depolarization (**Fig. 3**). As expected thermal acclimation strongly altered the threshold for cold-induced spreading depolarization (effect of acclimation: F_1,88_ = 72.4, P < 0.001) such that it occurred approximately 3.4°C lower (5.7 ± 0.4°C) in cold-acclimated flies than their warm-acclimated conspecifics (9.1 ± 0.4°C); a very similar difference was found in the sham-injected groups (5.8 ± 0.4°C and 9.1 ± 0.4°C for cold- and warm-acclimated flies, respectively) (**Fig. 3A**). Blockade of voltage-gated K^+^ channels and K^+^ channels in general led to a marked increase in the spreading depolarization temperature (effect of treatment: F_3,88_ = 14.1, P < 0.001), however, this effect was notably larger in cold-acclimated flies (interaction: F_3,88_ = 5.5, P = 0.002). Specifically, cold-induced spreading depolarization in cold-acclimated flies injected with 4-AP and TEA happened at 8.3 ± 0.5°C and 8.8 ± 0.4°C, respectively, while it happened at 10.2 ± 0.3 °C and 9.5 ± 0.4°C in warm-acclimated flies, reducing the effect of acclimation on the event from approximately 0.34°C per °C change in acclimation temperature to ∼ 0.13°C per °C (based on means of blocked and non-blocked groups; **Fig. 3A**).

**Figure 3.**
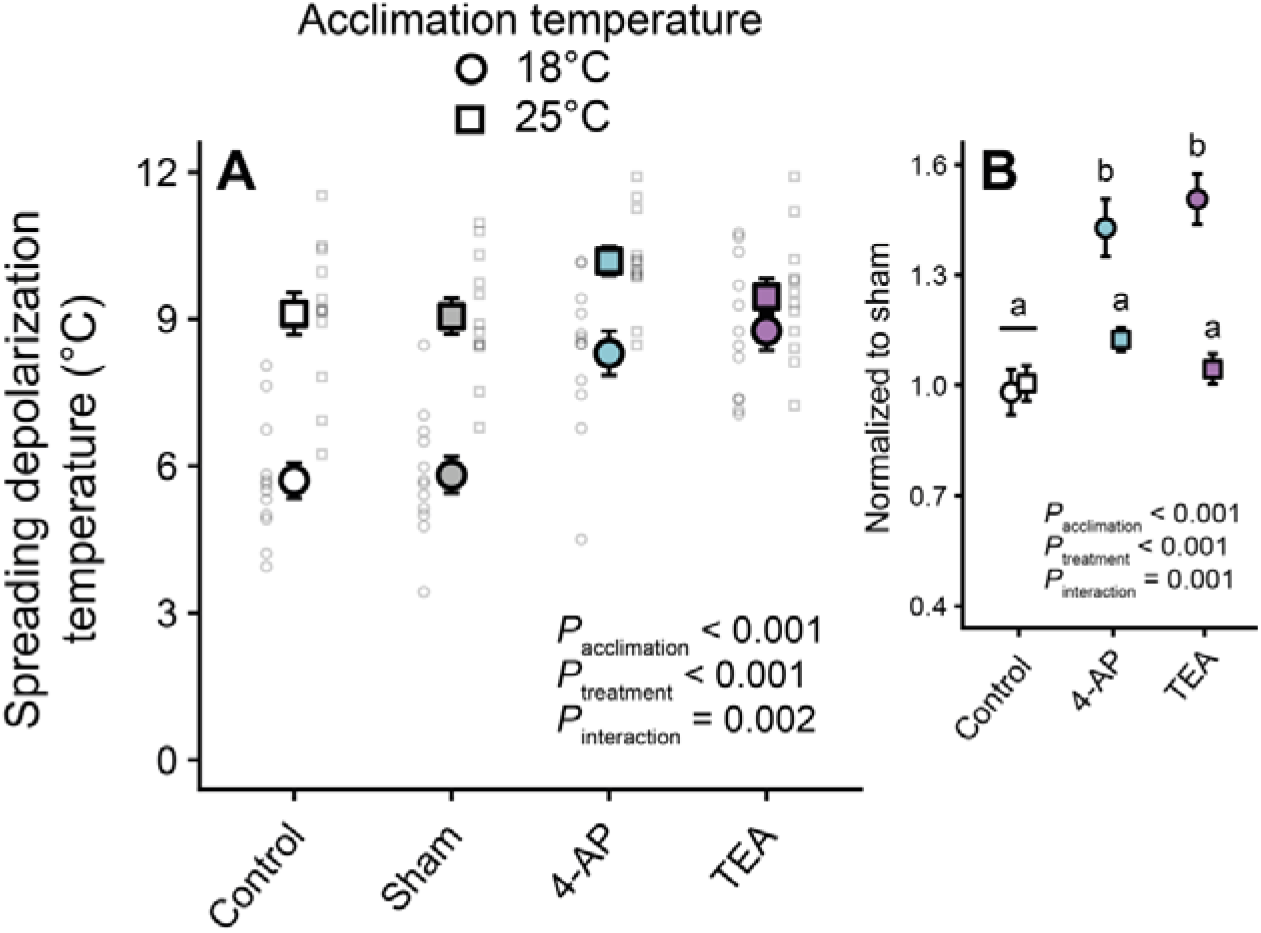
– Pharmacological blockade of K^+^ channels increases the threshold for cold-induced spreading depolarization in cold-acclimated flies. (A) The temperature leading to a cold-induced spreading depolarization differed between cold- and warm-acclimated flies (circles and squares, respectively), and was not affected by a sham injection. The addition of a voltage-gated K^+^ channel blocker (4-aminopyridine, 4-AP; blue) and a general K^+^ channel blocker (tetraethylammonium, TEA; purple) increased the temperature leading to spreading depolarization. (B) By normalizing to the respective sham-injected groups, the isolated effect of the pharmacological interventions revealed that the block of K^+^ channels was more pronounced in cold-acclimated flies, and that the block of voltage-gated K^+^ channels explained a large proportion of the general K^+^ channel block. Small transparent points in panel A represent individual data points for each group with data points for cold- and warm-acclimated flies being depicted on the left (small circles) and right (small squares) of group means, respectively. N = 12 flies per acclimation and treatment group combination; total 96 flies.

After normalization of each acclimation group to their respective sham groups (**Fig. 3B**), acclimation played a major role in determining the magnitude of change induced by K^+^ channel inhibition (effect of acclimation: F_1,66_ = 20.6, P < 0.001). Similarly, clear differences were found between the three experimental groups (i.e. control, 4-AP, or TEA; effect of treatment: F_2,66_ = 16.0, P < 0.001). These effects were largely driven by the magnitude of the response to inhibition in the cold-acclimated flies (interaction: F_2,66_ = 8.1, P = 0.001), where 42.8 ± 7.7% and 50.7 ± 6.8% increases were observed for 4-AP and TEA, respectively, whereas no effects were observed in warm-acclimated flies (**Fig. 3B**). From these observations, we can gather that the effect of voltage-gated K^+^ channel block represented a large proportion (∼ 84%) of the effect of the general K^+^ channel block in cold-acclimated flies (group estimates were similar based on a Tukey’s HSD *post hoc* test, P = 0.950).

### RNAi screen of voltage-gated K^+^ channel-encoding genes

After confirming the role of voltage-gated K^+^ channels in setting the threshold for cold-induced spreading depolarization in *D. melanogaster* we used the UAS/Gal4 model system to investigate which voltage-gated channels were likely to be involved. Specifically, we used a glial driver line (*repo*-Gal4) and TRiP RNAi lines (UAS-TRiP-gene^RNAi^) to knock down a range of channel-encoding genes and test how this impacted the temperature leading to cold-induced spreading depolarization (**Fig. 4**). Glia were targeted as spreading depolarization starts here in insects (9, 72). Here we found significant effects of knockdown on the temperature of spreading depolarization for three genes: *eag*, *sei*, and *Shaw* (P = 0.019, P = 0.001, and P = 0.001, respectively; see **Table S1** for all statistical outputs). In the case of the *eag* knockdown, however, the spreading depolarization temperature did not differ from all three control crosses and we therefore interpreted this as a non-significant result. Glial knockdown of channels encoded by *sei* and *Shaw*, by contrast, increased the temperature leading to cold-induced spreading depolarization by mean values of 1.67°C and 1.66°C, respectively (mean based on all control crosses).

**Figure 4.**
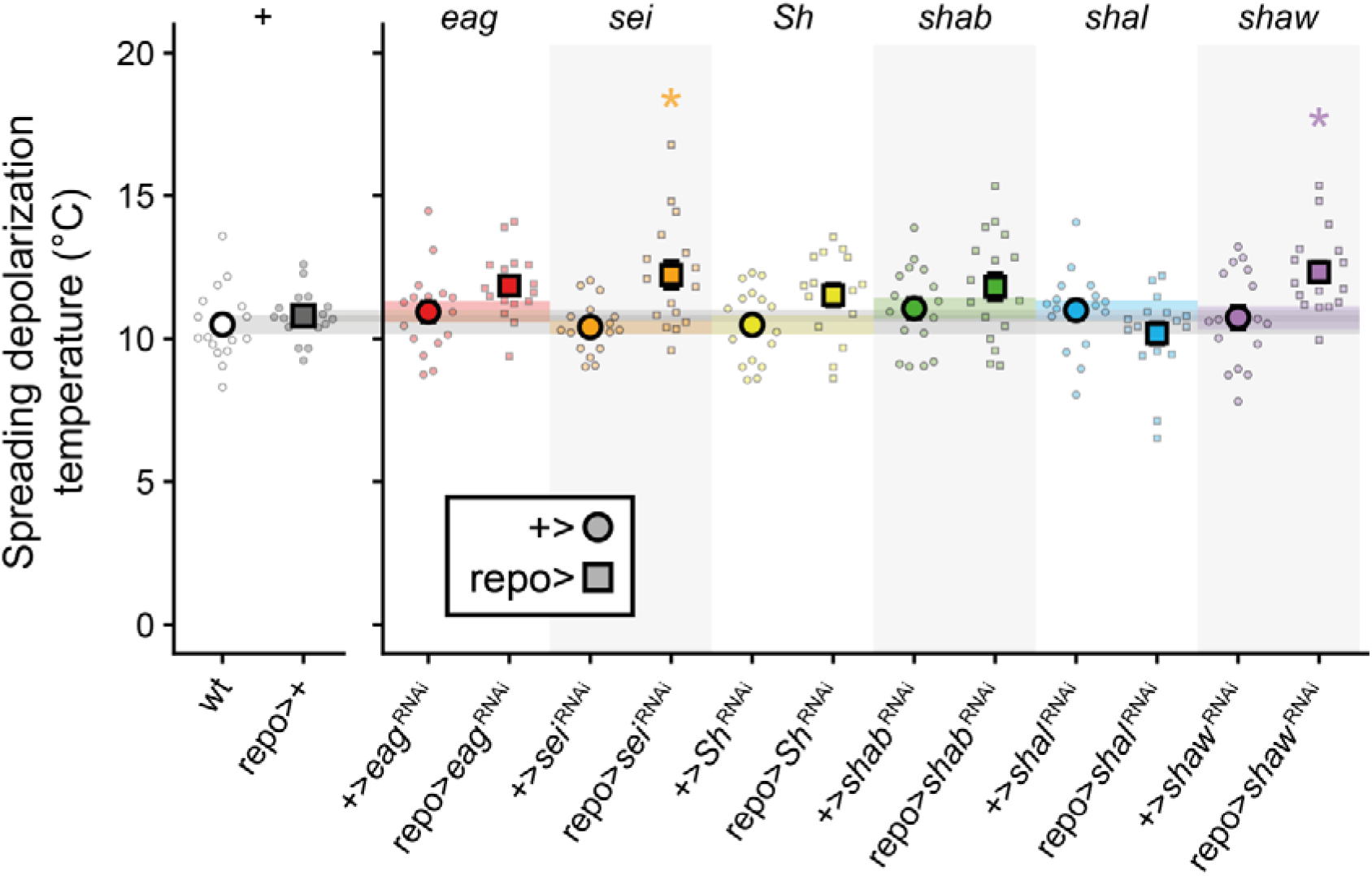
**– Targeted RNAi knockdown of glial *seizure* and *Shaw* reduces cold tolerance in *Drosophila melanogaster.*** A screen of RNAi-induced knockdown of channel-encoding genes, using the UAS/Gal4 model system and targeted at the glia (*repo*-driver), was used to identify potential molecular mechanisms underlying variation in the temperature leading to cold-induced spreading depolarization. In the left panel are two control crosses (wildtype [white circle] and driver [grey square] controls), while each gene targeted (right panel) has an extra control cross (reporter control [coloured circle]) and the RNAi-mediated knockdown [coloured square]). For each cross, individual data points are jittered around a large, opaque mean value point with the associated standard error bar (which is often obscured by the point representing the mean). Shaded horizontal panels in grey represent the standard error for each of the control crosses while those in colour represent that of each genes third control cross. Asterisks indicate where a statistically significant effect of the knockdown was found (see *Materials and Methods* for analysis). See **Table 1** for sample sizes.

*The effect of glial* sei *knockdown on anoxia and heat resistance*

To determine whether the *sei* gene is involved in tolerance to another abiotic stressor where spreading depolarization has shown to be involved, we tested how knockdown of this gene in the glia affected resistance to anoxia (**Fig. 4**). Subsequently, we tested its role in heat resistance, which also served as a test to see whether our cross had similar effects to those obtained previously (69).

For resistance to anoxia we found clear differences among crosses (F_3,60_ = 6.3, P = 0.001), and these differences were driven by the knockdown of *sei* having a ∼ 1.9 s shorter time to anoxia-induced spreading depolarization compared to the three control crosses, for which it occurred at 14- 15 s (**Fig. 4A**; see **Table S2** for all statistical outputs). The time to recover from anoxia upon the reintroduction of oxygen ranged from ∼ 12-24 s and was unaffected by *sei* knockdown (F_3,59_ = 1.3, P = 0.299; **Fig. 4B**).

Heat resistance was measured on the organismal level to repeat the procedure of a previously published study using a similar *sei* knockdown (69), except we noted down the time to both loss of coordination and complete paralysis (**Fig. 5**). For both phenotypes we found clear differences between crosses (loss of coordination: F_3,40_ = 7.9 with P < 0.001, and paralysis: F_3,40_ = 11.4, P < 0.001 – see **Table S2** for all statistical outputs), which was driven by the *sei* knockdown cross experiencing loss of coordinated movements on average 1.6 min earlier than control groups (∼ 4.1 min compared to 5.5 - 5.7 min, **Fig. 5A**) and complete paralysis ∼ 1.5 min earlier (5.4 min compared to 6.6 – 7.2 min, **Fig. 5B**).

**Figure 5.**
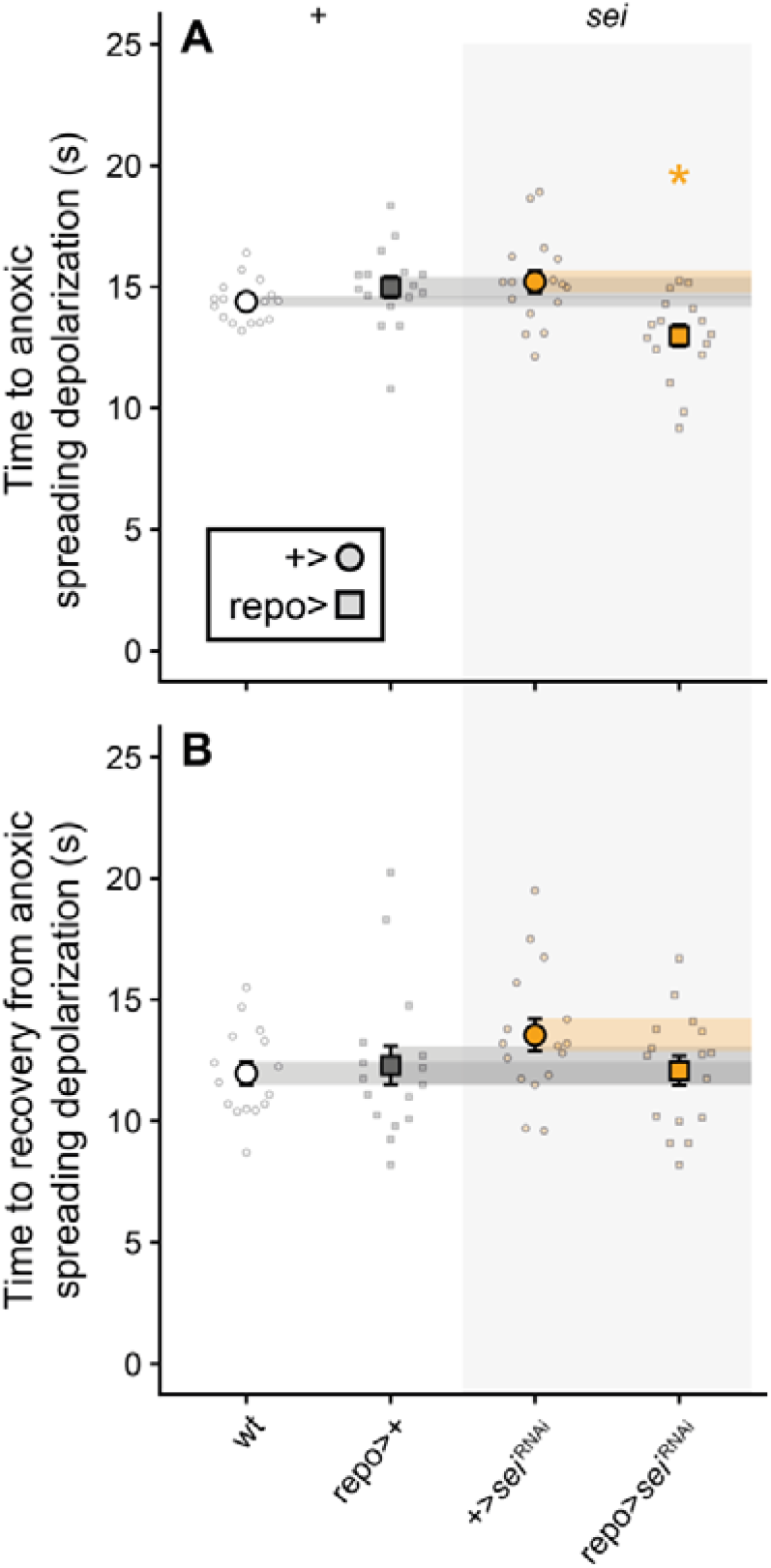
– Glial RNAi knockdown of a voltage-gated K^+^ channel encoded by the *seizure* gene reduces central nervous system resistance to anoxia with no effect on recovery. (A) RNAi- mediated knockdown of the *sei* gene, achieved with the UAS/Gal4 model system, revealed a role for channels encoded by *sei* in the ability to resist spreading depolarization when exposed to an atmosphere of pure N_2_. (B) Time to recovery from SD after the reintroduction of oxygen was unaffected. Crosses are as follows (left to right): wild type, driver control, reporter control, and knockdown. Individual data points are jittered around their respective means and standard errors. Error bars not shown are obscured by the mean symbols. Shaded panels across the figure represent the standard error for each of the control crosses. See **Table 1** for sample sizes.

## Discussion

In this study, we have shown for the first time that glial voltage-gated K^+^ channels promote cold acclimation of the CNS, and linked tissue-specific expression of channel-encoding genes (*sei* and to a lesser extent *Shaw*) to the organismal stress tolerance phenotypes through their involvement in the well-established neurophysiological phenomenon of spreading depolarization (SD). Specifically, we reaffirm a strong thermal acclimation response the *D. melanogaster* CNS and used a pharmacological approach to demonstrate that K^+^ channels, specifically the voltage-gated subtypes, play a key role in this process. From here, we screened for candidate voltage-gated K^+^ channels by using the UAS/Gal4 model system to knock down channel-encoding genes in glial cells and found that knockdown of *sei*- and *Shaw*-related channels reduced resistance to cold-induced SD similar to the pharmacological blockade of K^+^ channels in cold-acclimated flies. Lastly, we demonstrate a role for *sei*-related K^+^ channels in the general neural stress response as its knockdown reduced tolerance to anoxic SD (without affecting recovery) and heat-induced paralytic and comatose phenotypes.

Thus, while SD represents a clear limit to organismal function under severely unfavourable abiotic conditions (i.e. thermal or anoxic coma phenotypes (11, 12, 14)), the amount of stress needed to elicit SD can differ greatly, even within species, and this variation can be induced by differential expression of single genes.

### Pharmacological blockade of K^+^ channels eliminates the resistance to cold-induced spreading depolarization elicited by cold acclimation

To test whether K^+^ channels promote resistance to cold-induced SD, we compared the effects of pharmacological blockers in cold- and warm-acclimated *D. melanogaster*. Like previous studies, we found a large and pervasive effect of acclimation on the temperature leading to cold-induced SD in *D. melanogaster* (∼ 0.45°C per 1°C difference in acclimation temperature) (33, 36, 45). This acclimation effect was unaffected by sham injection of a simple physiological saline (**Fig. 1**). By contrast, application of 4-AP (voltage-gated K^+^ channel blocker) or TEA (general K^+^ channel blocker) both increased the low-temperature threshold for SD initiation of the cold-acclimated flies, reducing their cold resistance, and no observable effect was seen in the warm-acclimated conspecifics. This effect was biggest with TEA, however, the blockade of voltage-gated K^+^ channels with 4-AP made up a considerable proportion of the effect of broadly blocking K^+^ channels (**Fig. 3B**). This suggests that voltage-gated K^+^ channels are more heavily involved in improving CNS resistance to cold than other K^+^ channels. This is the first time voltage-gated K^+^ channels have been implicated in long-term acclimation effects on cold-induced SD. Previous studies on SD in locust CNS have shown protective effects of K^+^ channel inhibition on ouabain-induced SD and that K^+^ channel block can confer resilience to anoxia-induced SD in a manner similar to rapid cold hardening (16, 47, 61, 85). These findings contrast our findings of reduced tolerance with K^+^ channel inhibition; however, there are several methodological and experimental differences between our studies, most notably the drug dosage. Regardless, the authors of the locust studies argued that the positive effects of K^+^ channel block might be due to reduced neuronal release of K^+^ into the interstitium (16, 61). However, given the role of K^+^ channels in maintaining the interstitial K^+^ concentration *via* glial spatial buffering one could also predict a negative impact, as seen in rat brain slices which become more susceptible to SD with 4-AP or TEA application (62). Without further evidence, it seems most likely that these differences arise due to different mechanisms for short- vs. long-term cold pre-treatment (i.e. hardening vs. acclimation) or due to dosage differences (as the previous studies found protective effects of TEA independent of pre-treatment). Regardless, we expect that cold-acclimated fruit flies have improved resilience to cold-induced SD due to improved K^+^ buffering capacity, as previously suggested (33). This effect could be driven by increased abundance of membrane-bound voltage-gated K^+^ channels, which is counteracted by the application of K^+^ channel blockers. Alternatively, increased resistance to SD could be conferred by increased resistance to cellular depolarization through increased abundance of voltage-gated K^+^ channels, which open in response to cell depolarization to increase K^+^ permeability and stabilize the membrane potential, as seen in locust muscle fibres (86, 87). The remaining difference between voltage-gated- and general K^+^ channel block should be the non-voltage-gated channels, of which ATP-sensitive, Ca^2+^-activated K^+^ channels are known to be involved (63, 64). Similarly, it is well-established that changes to Na^+^/K^+^-ATPase activity and thermal sensitivity are involved (45, 46, 88). More research is needed to understand the role of K^+^ channels in the abiotic stress response of the CNS and how they modulate the susceptibility to stress-induced SD.

### Glial sei and Shaw knockdown reduce resistance to cold-induced spreading depolarization

After establishing a key role of voltage-gated K^+^ channels in CNS resilience to cold-induced SD, we sought to investigate which specific channels were responsible. We did so by using the UAS/Gal4 transgenic model system to knock down the expression of voltage-gated K^+^ channel-encoding genes, and we decided to drive this expression in the glia due their well-documented role in spatial buffering and neuroprotection, and recent evidence that SD is triggered at the glial hemolymph-brain barrier in insects (70–72). From these experiments, we found that the glial knockdown of channels encoded by the genes *sei* and *Shaw* increased the temperature leading to cold-induced SD (**Fig. 4**). All other knockdowns, except *Shal*, showed the same trend. Thus it seems likely that the improved resistance to cold-induced SD after cold acclimation was due to increased glial expression of voltage-gated K^+^ channels encoded by these genes.

*sei* encodes an inward rectifying channel ortholog to the human hERG channel (K_v_11.1) and responds to mild depolarization by opening and promoting inward K^+^ movement away from the extracellular space (89). *Shaw* on the other hand encodes a delayed, non-inactivating rectifier channel ortholog to the human K_v_3 family that is only mildly voltage-sensitive and therefore thought to function as a leak channel (90, 91). Thus, the knockdown of genes encoding channels responsible for facilitating K^+^ currents in opposite directions lead to the same phenotype. The mechanism underlying this remains unknown, however, we speculate that both channels play a role in glial spatial buffering which requires bidirectional K^+^ movement (58), and future studies should aim to elucidate the potential cooperative role of *sei*- and *Shaw*-related channels in this process.

Lastly, it is interesting to note that both *sei* and *Shaw* are expressed in both neurons and glia, with expression in glia being substantially lower than that in neurons (92, 93). Thus, while we targeted the glia for knockdown in this study, it is possible that *sei* and *Shaw*- related channels play different roles in the neurons, which could interact with glial spatial buffering mechanisms. Regardless, the effect of the glial knockdown here indicates a particularly critical role in the glia.

This is not the first time *sei* channels have been implicated in the response of neural tissue to abiotic stress; loss of *sei* expression or function reduced organismal heat tolerance of *D. melanogaster* (69), indicating that these channels are involved in a general thermal stress response of the CNS. Furthermore, the authors showed that UAS/Gal4 knockdown targeted at both neurons and glia reduce tolerance, and further demonstrated that the loss of *sei* expression in octopaminergic neurons, in particular, drives this response. Octopamine is a strong modulator of glial K^+^ conductance (94), and the octopaminergic pathway has been highlighted as a mechanism of improved resilience to SD in rapid cold hardening (48). Indeed, *sei* and octopamine receptor genes show a large degree of co-expression in *D. melanogaster* neurons (95). It is therefore likely that the changes in K^+^ conductance elicited by octopamine occur through *sei*-related channels and by extension that their co-expression is essential for promoting insect CNS cold, or stress, tolerance.

We did not investigate neuronal expression of *sei*, however, our results highlight the importance of *sei*-related channels in glial cells in the thermal stress response, and we speculate that this is due to a yet-to-be-investigated role in glial spatial buffering in the insect CNS. The *repo*-driver is a pan-glial driver and its use in knockdowns therefore provides little evidence regarding the subtype involved in the phenotype studied. However, the effect of pan-glial *sei* knockdown on CNS heat stress tolerance is known to be elicited by knockdown in neuropil-ensheathing and perineurial glia (69). Perineurial glia are involved in the barrier function of the glial hemolymph-brain barrier, while the neuropile-ensheathing glia are integral in forming an internal diffusion barrier that protects the neuropil within the ganglion (96, 97). Thus, *sei* is integral in K^+^ diffusion barrier formation in polarized brain barriers. As *sei* encodes an inwards rectifier, one would expect its expression to be adglial (facing the neuropil) if K^+^ barrier function is to be maintained, which would also promote removal of K^+^ from the interstitium and glial K^+^ buffering. Consequently, we argue that glial K^+^ spatial buffering is essential to reduce susceptibility to SD and that channels encoded by *sei* are integral to this process.

The *Shaw* gene encodes a presumed outward rectifying leak channel, and its knockdown also elicited reduced cold tolerance of the CNS (i.e. SD at a higher temperature, **Fig. 4**). In the current paradigm of SD susceptibility, greater leak of K^+^ into the interstitium is considered maladaptive (10, 16) and we were therefore surprised to see that the knockdown of this K^+^ leak channel in glia reduced cold stress tolerance of the CNS. The cellular or glia-subtype-specific localization of *Shaw* has not been investigated; however, should *Shaw*-channels have a role in the glia (i.e. K^+^ buffering) one could speculate that it is expressed in the glial membrane facing away from the neurons (i.e. basolateral) to promote spatial K^+^ buffering (58). Interestingly, there is a continuous flux of K^+^ out of the ganglia of insects which is assumed to occur through leak channels (98). Thus, it is compelling to argue that *Shaw* channels expressed in perineurial glia facilitate this outward K^+^ flux, and by extension that their knockdown prevents proper removal of K^+^ from the ganglion and its interstitium, resulting in impaired spatial buffering and SD susceptibility.

An alternative role for outwards-rectifiers like *Shaw*-channels, could be their ability to improve maintenance of membrane polarization, as their opening will act to repolarize a depolarized membrane potential by increasing K^+^ permeability, as seen in muscle fibres of cold-acclimated locusts (86). Thus, if the glial membrane potential and its depolarization were part of the SD initiation mechanism, reducing K^+^ permeability might reduce resistance to SD, which is what we observe here. In the locust muscle, the depolarization resistance was achieved through increased expression of *Sh*-related channels; however, we found no effect of glial *Sh* knockdown on SD susceptibility (**Fig. 4**), and as such muscle cells and glia may rely on modulating different K^+^ currents or channels to facilitate tissue-specific cold acclimation. Regardless, more research is needed to identify causal links between K^+^ channels abundance and localization, glial spatial buffering, and SD.

### Glial sei knockdown reduces the general stress tolerance of the central nervous system

After establishing a role for *sei* in modulating cold-induced SD, we wanted to test whether this was a general mechanism that can induce variation in SD sensitivity in other abiotic stressors (11, 12, 16). We tested how glial *sei* knockdown affected the ability to resist anoxia-induced SD and found that it increased SD susceptibility by reducing the time to anoxic SD without any effect on recovery times (**Fig. 5**). Similarly, we repeated the experiments of Hill et al. (69) and found that the knockdown of *sei* in glia cells reduced organismal heat tolerance as *sei* knockdown flies became paralytic and comatose sooner during heat exposure (**Fig. 6**). Bear in mind that while we didn’t measure heat-induced SD, the link between heat-induced paralysis and SD is well-established in *Drosophila* (11). Thus, our results effectively link tissue-specific expression of a single gene to general, organismal stress tolerance through the loss of CNS function by SD.

**Figure 6.**
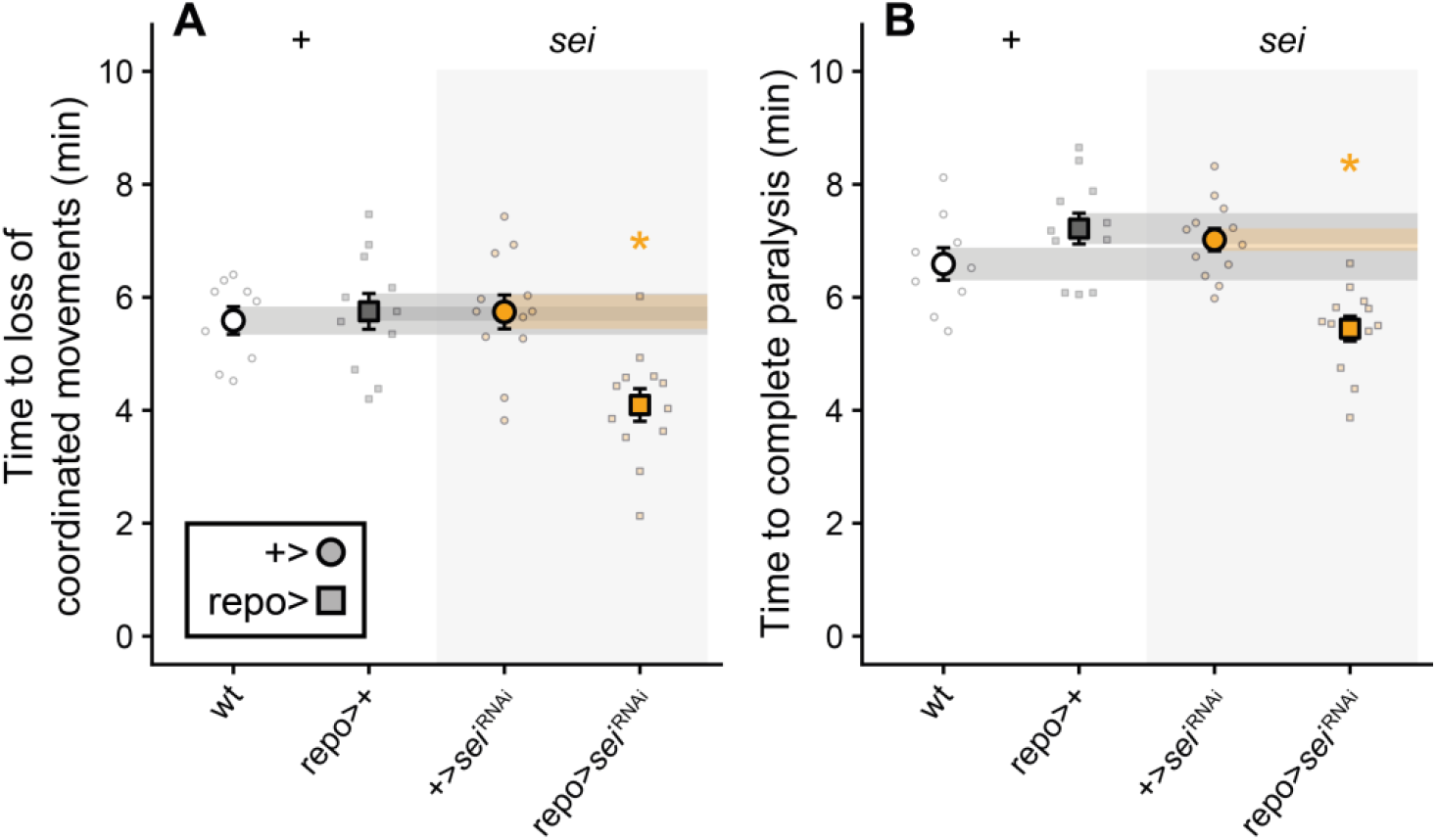
– RNAi-mediated knockdown of the *seizure* gene in glia reduces organismal heat tolerance. Time to loss of function was measured as the time to (A) loss of coordinated movements and (B) complete loss of function. Overall, the glial knockdown of the *sei* gene resulted in a reduction in heat tolerance, similar to how cold- and anoxia-tolerance were impacted. In each panel, crosses are as follows (left to right): wild type, driver control, reporter control, and knockdown. Individual data points are jittered around their respective means and standard errors. Error bars not shown are obscured by the mean symbols. Shaded panels across the figure represent the standard error for each of the control crosses. See **Table 1** for sample sizes.

Anoxia has been used to study the mechanisms of SD in insects since its initial discovery due to its ease of application at constant ambient temperature, as opposed to cold-or heat-induced SD where temperature-effects need to be considered (32, 44). Furthermore, anoxia has the benefit of immediately impairing ATP production in small insects (1). This means that for species like *D. melanogaster*, anoxia exposure tests the ability to resist SD in the absence of active transport, that is, the ability for passive mechanisms to resist SD (see discussion in (33)). Passive ion movement through voltage-gated ion channels does not require ATP, meaning that our findings of reduced resistance to anoxia-induced SD (**Fig. 3A**) provide additional support for channels encoded by *sei* being involved in the passive component of glial spatial buffering (i.e. through K^+^ channels independent of the ATP-dependent Na^+^/K^+^-ATPase) (58). Alternative passive mechanisms include, but are not limited to, decreased cellular leakage of K^+^, larger interstitial volumes (to dilute the increased K^+^ concentration during SD), or a combination of both. Changes to anaerobic ATP production also cannot be ruled out as a mechanism (99). However, it seems unlikely that glial knockdown of the inward rectifying *sei*-related channels would promote any of these processes.

Recovery from SD involves restoration of perturbed ion gradients and membrane potentials, and by extension neuronal excitability (9, 20, 72). Given the importance of glial ion regulation, and by extension the proposed role of glial *sei*-channels in spatial K^+^ buffering, in this process we were surprised to find that their knockdown had no effect on the time to recovery from anoxia-induced SD (**Fig. 5B**). That said, voltage-gated channels open during SD and tend to stay open for the duration of the SD, meaning further opening is unlikely to occur and therefore cannot aid during recovery (10, 100). Instead recovery is achieved *via* active transport by electrogenic ATPases, like the Na^+^/K^+^-ATPase and V-type H^+^-ATPase, which re-establish ion gradients upon return of ATP following reoxygenation (16, 72). Based on our findings, we therefore posit that the passive component of glial spatial buffering, of which *sei*-channels play a pivotal role, primarily promotes resistance to SD in the initial stress response of the CNS, while active ion transport mechanisms dictate the rate of recovery.

The purpose for testing the heat tolerance of glial *sei* knockdown flies was two-fold: It tested the repeatability of the *sei* knockdown (as compared to Hill et al. (69)), while also testing for the general involvement of *sei* channels in SD-related phenotypes (i.e. heat paralysis and coma). Thus, if our results match earlier reports we would expect the cross to a similar molecular phenotype of reduced channel abundance. Our findings closely match those of Hill et al., affirming not only the negative effects of glial *sei* knockdown on heat tolerance but also the efficacy of our crosses based on a different background line (i.e. Canton S vs. *w*^1118^). Similarly, our findings support the involvement of *sei* channels in the general susceptibility to abiotic stress-induced SD. Heat stress is generally thought to confer SD susceptibility through increased K^+^ leakage into the interstitium (16). Thus, in light of their hypothesized role in glial K^+^ buffering, presumably at the inward-facing membranes of perineurial or ensheathing glia (see discussion above), it seems likely that the knockdown of inward rectifying *sei* channels in the glia should lead to a reducing resistance to heat coma.

## Conclusion

In summary, we show that the improved cold resistance of the CNS conferred by cold acclimation is driven, at least partially, by expression of voltage-gated K^+^ channels. We further demonstrate that these channels are likely to be *sei*- or *Shaw*-related, and that channels encoded by the *sei* gene are responsible for modulating general susceptibility to stress-induced SD. Thus, we effectively link tissue-specific (glial) expression of a channel-encoding gene to its involvement in modulating the susceptibility to well-established neurophysiological phenomenon (SD) and ultimately to an organismal stress tolerance phenotype (coma). Such an important role for a voltage-gated channel in non-excitable glia seems unexpected, but falls in line with broader interest in the important roles voltage-gated channels play in non-excitable tissues (101). Nonetheless, further research is needed to establish the exact, causal mechanisms by which glial voltage-gated K^+^ channels modulate susceptibility to SD under stress. Future studies should focus on investigating the cellular and tissue-specific expression, abundance, and localization of voltage-gated K^+^ channels, particularly those encoded by the *sei* gene, and elucidating the processes by which interstitial K^+^ concentrations are managed during exposure to severe abiotic stress. Doing so will inevitably increase our understanding of the mechanisms that limit insect thermal tolerance, while also providing valuable, translational insights into the physiological mechanisms of human and mammalian neuropathologies associated with SD, like migraine and stroke.

## Acknowledgements

We would like to extend our gratitude to Dr. Woo-Jae Kim (University of Ottawa) for providing us with the Canton S and *repo*-Gal4 lines used in our experiments.

## Data availability

All data was made to reviewers during the review phase and can be found here (https://figshare.com/articles/dataset/Dataset_-_Andersen_Livingston_Robertson_MacMillan_-_Glial_voltage-gated_K_sup_sup_channels_modulate_the_neural_abiotic_stress_tolerance_of_i_Drosophila_melanogaster_i_/26124634 ).

## Author contributions

The study was conceived and designed by M.K.A., R.M.R., and H.A.M.. M.K.A. and D.B.H.L. performed the experiments, and M.K.A. analyzed the data, and wrote the first draft. All authors made edits to the manuscript and approved the final version of the manuscript.

## Funding

This research was funded by Carlsberg Foundation Internationalization Fellowships (CF18-0940 and CF19-0472) to MKA and a Discovery Grant from the Natural Sciences and Engineering Research Council of Canada (RGPIN-2018-05322) to HAM. Additional funding for this project came through grants to the TRIA-FoR Project (https://tria-for.ualberta.ca/) to H.A.M. from Genome Canada (project no. 18202), the Government of Alberta through Genome Alberta (project ID L20TF), and the Ontario Research Fund – Ontario Ministry of Colleges and Universities through Ontario Genomics (File No. 18202), with contributions from the University of Alberta, Carleton University, and the Great Lakes Forestry Centre Natural Resources Canada.

## Conflict of interest

None

## Supplementary material

**Table S1.**
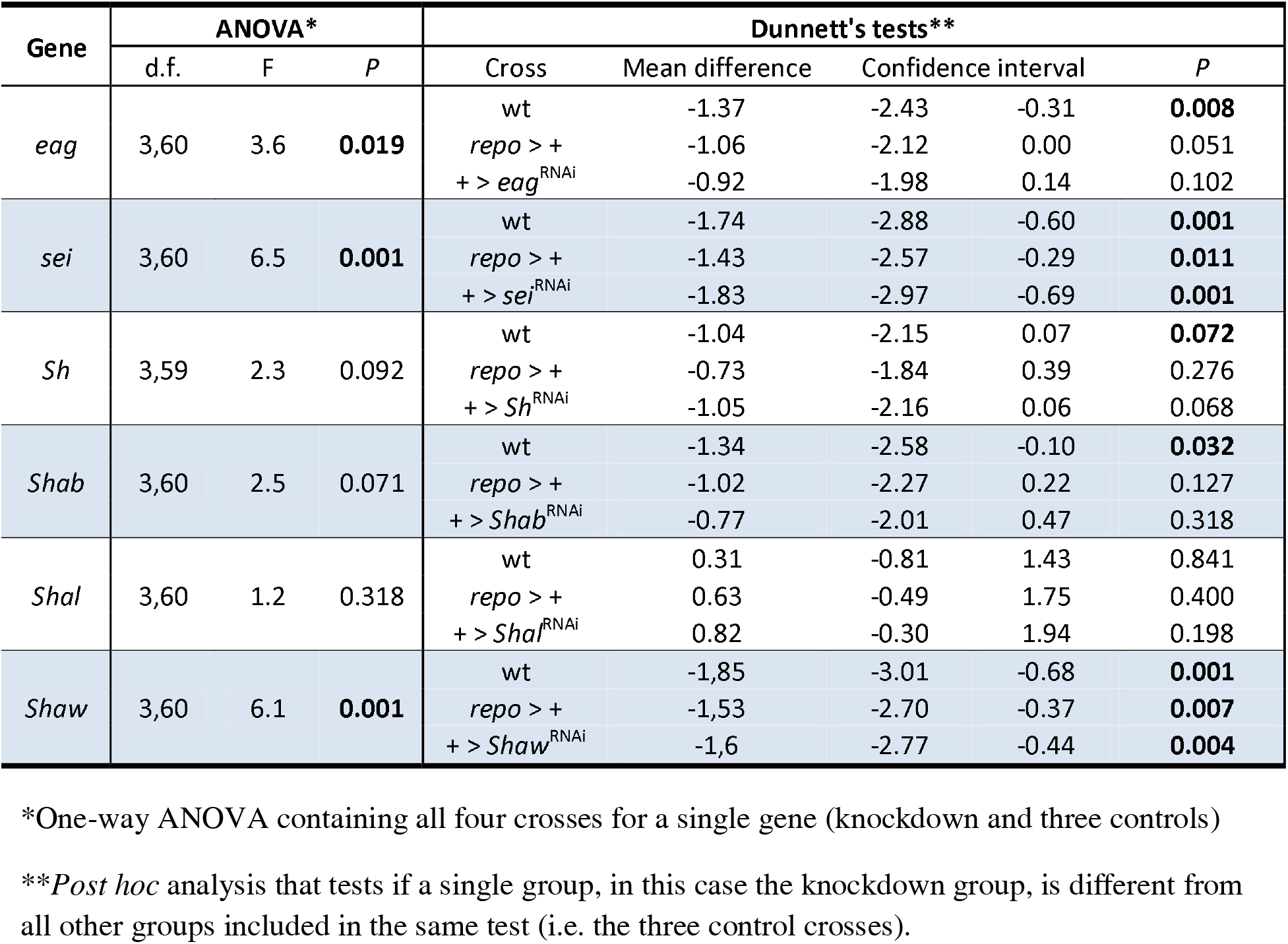
–. Output from statistical tests run on the UAS/Gal4-mediated RNAi knockdown of voltage-gated K^+^ channel-encoding genes in glia. The mean difference and confidence interval are in °C.

| Gene | ANOVA* |  |  | Dunnett's tests** |  |  |  |
| --- | --- | --- | --- | --- | --- | --- | --- |
|  | d.f. | F | P | Cross | Mean difference | Confidence interval | P |
| *eag* | 3,60 | 3.6 | **0.019** | wt | -1.37 | -2.43 -0.31 | **0.008** |
|  |  |  |  | *repo* > + | -1.06 | -2.12 0.00 | 0.051 |
|  |  |  |  | + > *eag*<sup>RNAi</sup> | -0.92 | -1.98 0.14 | 0.102 |
| *sei* | 3,60 | 6.5 | **0.001** | wt | -1.74 | -2.88 -0.60 | **0.001** |
|  |  |  |  | *repo* > + | -1.43 | -2.57 -0.29 | **0.011** |
|  |  |  |  | + > *sei*<sup>RNAi</sup> | -1.83 | -2.97 -0.69 | **0.001** |
| *Sh* | 3,59 | 2.3 | 0.092 | wt | -1.04 | -2.15 0.07 | **0.072** |
|  |  |  |  | *repo* > + | -0.73 | -1.84 0.39 | 0.276 |
|  |  |  |  | + > *Sh*<sup>RNAi</sup> | -1.05 | -2.16 0.06 | 0.068 |
| *Shab* | 3,60 | 2.5 | 0.071 | wt | -1.34 | -2.58 -0.10 | **0.032** |
|  |  |  |  | *repo* > + | -1.02 | -2.27 0.22 | 0.127 |
|  |  |  |  | + > *Shab*<sup>RNAi</sup> | -0.77 | -2.01 0.47 | 0.318 |
| *Shal* | 3,60 | 1.2 | 0.318 | wt | 0.31 | -0.81 1.43 | 0.841 |
|  |  |  |  | *repo* > + | 0.63 | -0.49 1.75 | 0.400 |
|  |  |  |  | + > *Shal*<sup>RNAi</sup> | 0.82 | -0.30 1.94 | 0.198 |
| *Shaw* | 3,60 | 6.1 | **0.001** | wt | -1.85 | -3.01 -0.68 | **0.001** |
|  |  |  |  | *repo* > + | -1.53 | -2.70 -0.37 | **0.007** |
|  |  |  |  | + > *Shaw*<sup>RNAi</sup> | -1.6 | -2.77 -0.44 | **0.004** |

\*One-way ANOVA containing all four crosses for a single gene (knockdown and three controls)
\*\**Post hoc* analysis that tests if a single group, in this case the knockdown group, is different from all other groups included in the same test (i.e. the three control crosses).

**Table S2.**
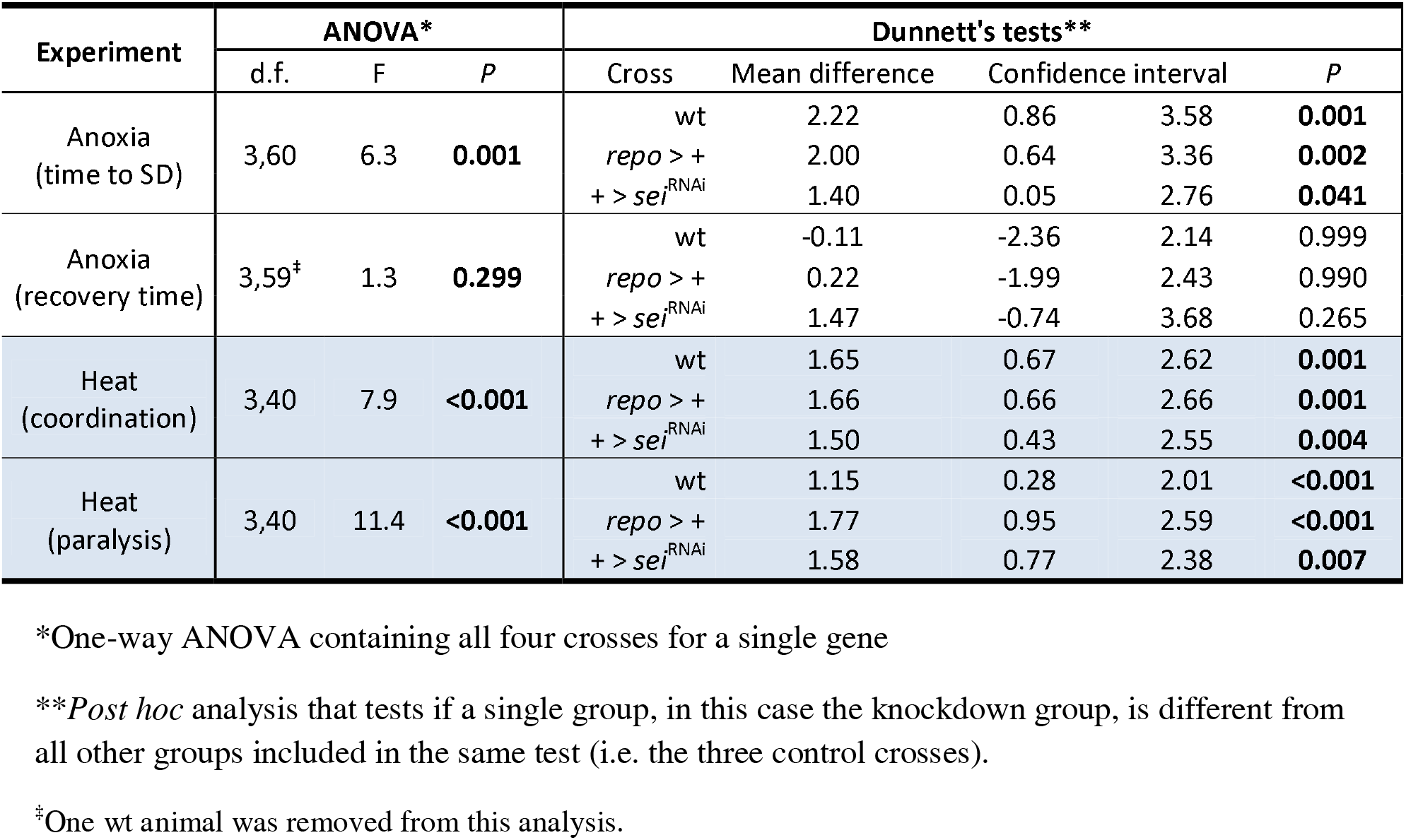
–. Statistical outputs for the analysis of the effect of *sei* knockdown on resistance to anoxia-induced spreading depolarization and heat-induced loss of organismal function. The mean difference and confidence interval are in s for anoxia resistance and in min for heat resistance.

## Notes

https://figshare.com/articles/dataset/Dataset_-_Andersen_Livingston_Robertson_MacMillan_-_Glial_voltage-gated_K_sup_sup_channels_modulate_the_neural_abiotic_stress_tolerance_of_i_Drosophila_melanogaster_i_/26124634

### Competing Interest Statement

The authors have declared no competing interest.

